# *In Silico* Processing of the Complete CRISPR-Cas Spacer Space for Identification of PAM Sequences

**DOI:** 10.1101/274670

**Authors:** Brian J Mendoza, Cong T Trinh

**Affiliations:** Department of Chemical and Biomolecular Engineering, University of Tennessee, Knoxville TN, USA

**Keywords:** PAM, protospacer, spacerome, CRISPR, Cas, CASPER, CASPERpam

## Abstract

Despite extensive exploration of the diversity of CRISPR (Clustered Regularly Interspaced Short Palindromic Repeats) systems, biological applications have been mostly confined to Class 2 systems, specifically the Cas9 and Cas12 (formerly Cpf1) single effector proteins. A key limitation of exploring and utilizing other CRISPR-Cas systems with unique functionalities, particularly Class I types and their multi-protein effector complex, is the knowledge of the system’s protospacer adjacent motif (PAM) sequence identity. In this work, we developed a systematic pipeline, named CASPERpam, that enables us to comprehensively assess the PAM sequences of all the available CRISPR-Cas systems in the NCBI database of bacterial genomes. The CASPERpam analysis revealed that within the 30,389 assemblies previously screen for CRISPR arrays, there exists 26,364 spacers that match somewhere in the viral, bacterial, and plasmid databases of NCBI, using the constraints of 95% sequence identity and 95% sequence coverage for blast hits. When grouping these results by species, we were able to identify putative PAM sequences for 1,049 among 1,493 unique species. The remaining species either have insufficient data or an undetermined result from the analysis. Finally, we were able to infer certain design principles that are relevant for understanding PAM diversity and a baseline for further experimental studies including PAM assays. We envision CASPERpam is a useful bioinformatic tool for understanding and harnessing the diversity of CRISPR systems.

## 1. Introduction

The bacterial adaptive immunity system CRISPR (Clustered Regularly Interspaced Short Palindromic Repeats), is near-ubiquitous across the archaea and bacterial kingdoms and exhibits extensive diversity [1]. Despite this heterogeneity, there exists two fundamental biochemical behaviors shared among all CRISPR systems: adaptation and interference [2]. The first stage, adaptation, is relatively conserved across species and involves the process by which foreign DNA (or RNA) is acquired and inserted into the CRISPR array. The second step, interference, involves the expression of the acquired sequences of the array and processing them into short RNAs for recognition and cleavage of the foreign DNA/RNA. To avoid self-cleavage, a short sequence known as a protospacer adjacent motif or PAM, appears just outside the recognition sequence. The PAM serves as a checkpoint to ensure the correct target has been found [3-5]. This first step is critical to the interference step, as even a perfectly matched guide RNA sequence without the proper PAM sequence will fail to achieve any activity [4, 6].

Significant research has been devoted to the classification of CRISPR-Cas systems to help define common functionalities [1, 7]. The current system contains two main classes, distinguished by a single (Class 2) or multi (Class 1) protein effector complex and further classified into 6 types and 22 subtypes [1]. Experimental studies have shown however that PAM sequences are particularly diverse and may vary from species to species even within subtypes [8]. The knowledge of the PAM sequence is thus critical for harnessing CRISPR-Cas systems, and the current classification system offers incomplete insight into a species’ PAM sequence. As research increasingly branches out from the canonical Cas9 of *Streptococcus pyogenes* and its Type II relatives to discover novel functionalities in alternate CRISPR-Cas systems, the ability to study these systems will be limited to the speed in which PAM sequences can be determined [9, 10].

Three common methods exist for determining the PAM sequence and have been used to varying degrees of efficacy: *in vivo* selection [11-13], *in vitro* assays [13, 14], and *in silico* analysis [4, 15, 16]. The first two methods are experimentally driven and provide the most accurate determination of the PAM sequence. However, they require time-intensive experimental procedures, and rely on effective molecular biology tools suited to the organism in question to be previously developed (*in vivo*), or the ability to clone out and express an active system heterologously (*in vitro*) [5]. In contrast, the *in silico* method uses sequence similarity of existing spacers in an organism’s genome to that of invasive genetic elements to suggest a potential PAM sequence [15]. While the method is less accurate and complete than that of its experimental counterparts, it requires simply a query of CRISPR array(s) against potential target genomes, which are readily available in ever-expanding sequencing databases. Although this method has been performed for a number of organisms to elucidate potential PAM sequences, a comprehensive PAM analysis of all available CRISPR-carrying organisms has not previously been performed.

We have recently presented the CRISPR Associated Software for Pathway Engineering and Research (CASPER) for enhanced on- and off-target prediction of gRNA designs and multi-targeting analysis for single and consortium of organisms using various Cas enzymes [17]. In this study, we have developed the CASPERpam algorithm to systematically and comprehensively perform a complete PAM analysis for all prokaryotes in the NCBI database for which sufficient data exists to investigate potential target matches. From the generated PAM sequence database, we were able to infer certain design principles that are relevant for understanding PAM diversity and provide a baseline for further experimental studies including PAM assays. The PAM sequence database and categories have been integrated into the CASPER platform as a new feature. We anticipate the CASPERpam tool will enable further research and engineering of prokaryotes using their native CRISPR machinery and aid in the development of novel CRISPR-Cas systems for heterologous expression in eukaryotes.

## 2. Methods

This section presents the CASPERpam algorithm for processing the complete CRISPR-Cas spacer space to identify PAM sequences to complement the resources of CASPER [17] for research in novel CRISPR-Cas systems. The CASPERpam algorithm contains 5 steps for spacer collection, genome collection, and PAM detection and analysis (Figure 1).

**Figure 1.**
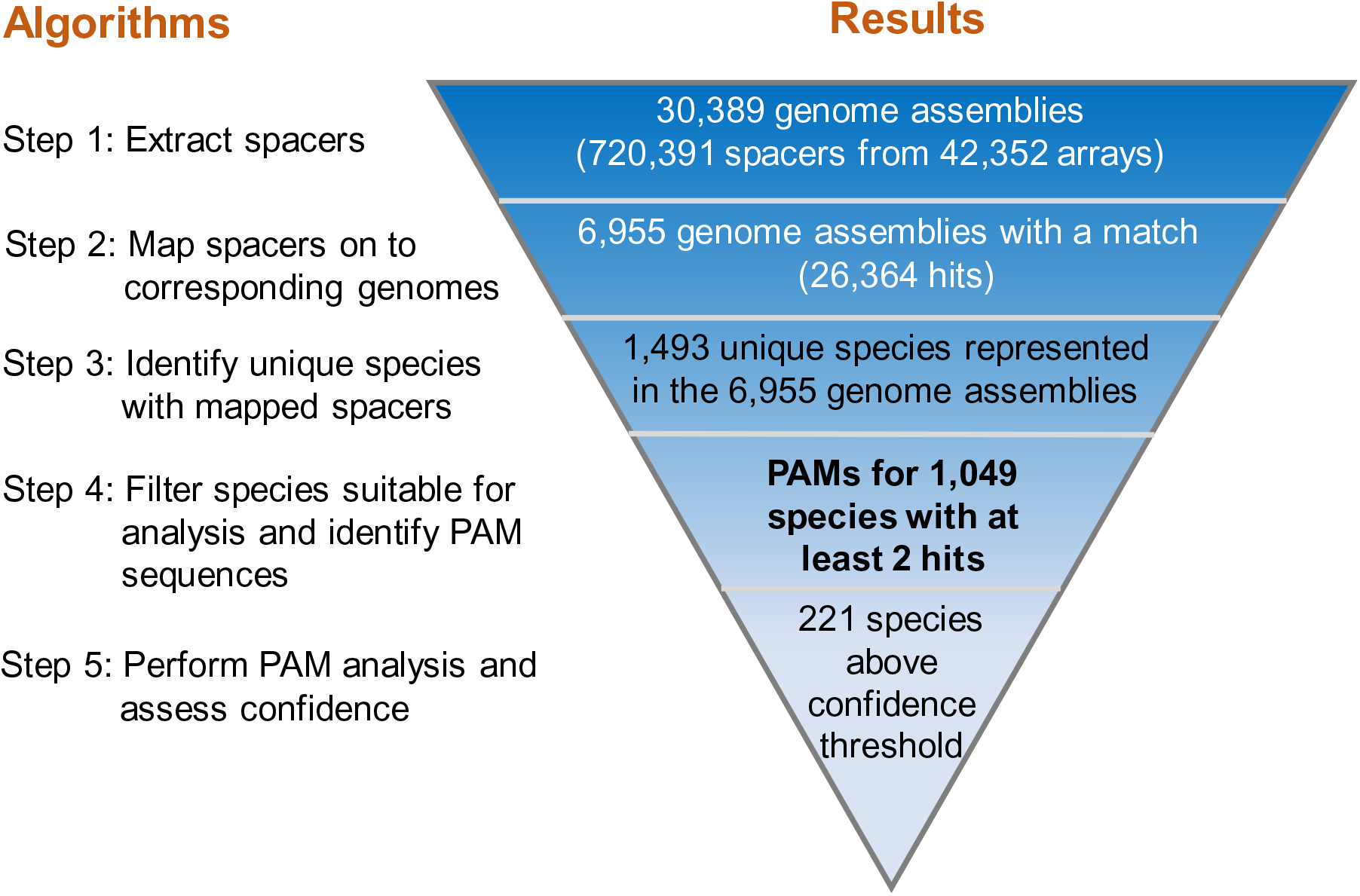
CASPERpam algorithm for identifying PAM sequences

### 2.1. Spacer collection for PAM analysis

Based on the 720,391 spacers detected across 42,352 CRISPR arrays from 30,389 assemblies collected and analyzed by Shmakov *et al.* [18], a total of 26,364 hits were detected by a BLASTN search across the viral (13,885 assemblies), plasmid (11,218 assemblies), and bacterial/prokaryotic (12,9209 assemblies) genome databases of NCBI, using the 95% sequence identity and 95% coverage constraints to gather only hits with sufficient similarity (Step 1, Figure 1). There were no spacers that generated multiple hits. To identify a PAM sequence, we collected the hit sequence and included an additional 10 nucleotides on either side (the “collected” sequence). Because the hit sequence itself may only be a partial match to the spacer, we extrapolated endpoints of the spacer within the collected sequence. Specifically, the spacer and collected sequences were aligned by finding the largest perfectly matched sequence shared between the two sequences. Then the indices of the spacer sequence’s start and end within the collected sequence were used to define the locations of the flanking sequences.

### 2.2. Genome collection for PAM analysis

From the spacer collection suitable for PAM analysis, we could narrow the number of assemblies from 30,389 to 6,955 (Step 2, Figure 1). Since multiple assemblies can belong to the same species, we clustered them at the species level to create a total of 1,493 species (Step 3, Figure 1). By choosing species with at least 2 spacer hits, we further reduced the number of species from 1,493 to 1,049 for which we could search for PAMs (Step 4, Figure 1). In our analysis, strains and substrains were grouped together under a common species. While in some rare cases it is possible that PAM sequences might differ between strains and substrains due to random mutation or perhaps horizontal gene transfer of CRISPR-Cas machinery among strains [7], we would not have been able to perform further analysis in a vast majority of species due to the limited number of hits belonging to each substrain.

### 2.3. PAM detection and analysis

The Cas type was used to determine whether a PAM was located 5’ (upstream) or 3’ (downstream) of the hit (Step 5, Figure 1). For type I, III (rPAM), IV, and V systems, the PAM has been either shown or inferred to occur 5’ of the spacer sequence [19-21]. Type II systems, however have 3’ PAMs of the spacer sequence. Each species was assigned a CRISPR-Cas system based on the data provided by Makarova *et al.* [1, 7]. For those species that did not appear in the prior classification studies, the genus was used to match the species to a relative and its Cas type was inferred from the type of its relative.

To obtain the PAM for each of the 1,049 species, bitwise scoring [22] was used to determine the important positions (Step 5, Figure 1). Positions with an R-score higher than a half a standard deviation above the mean were considered significant. As an additional constraint, the R-scores for positions farther than 5 nucleotides away from the spacer sequence were subject to the more stringent requirement of full standard deviation above the mean to further filter out noise from positions unlikely to contribute to the PAM sequence. Within each significant position, if a nucleotide had a relative frequency higher than 0.5, it was considered to be the consensus nucleotide. If no nucleotide was higher than 0.5, the nucleotide(s) that had relative frequency greater than 0.25 were considered to be part of the PAM.

The R tool ggseqlogo was used to create a bitwise graph for each species to determine the PAM from the appropriate flanking sequence (5’ or 3’) [22]. In many cases, sequences obtained were shorter than 10 nucleotides due to the spacer index starting in the flanking sequence and/or appearing at the end of an assembly or near an ambiguous sequence (set of ‘N’s). To satisfy the symmetrical sequence length requirement of ggseqlogo, flanking sequences smaller than 6 were discarded and the remaining were truncated to the length of the smallest sequence.

### 2.4. Computation

NCBI servers were used solely for query purposes (https://www.ncbi.nlm.nih.gov). All genomes needed to identify flanking sequences of the spacer hits were downloaded from NCBI (RefSeq/GenBank) onto an external hard drive (as of 11/24/2017). Python (ver. 3.6) was used to process the data and sequences. R (ver. 3.4.3) statistical programming language was used to run ggseqlogo and generate the sequence bitwise plots [22]. All computational work was performed on a Macintosh MacBook Pro computer (Apple Inc., 2013). Code is available at https://github.com/TrinhLab/CASPERpam.

## 3. Results and Discussion

### 3.1. Predictive PAM sequences match experimental evidence

To validate the CASPERpam algorithm, we compared the predicted and experimentally derived PAM sequences for a set of 12 species spanning both CRISPR classes and multiple types/subtypes (Figure 2). Our result shows that CASPERpam could generate the exact match (green boxes) for the experimentally derived PAM sequences in 3 of the 12 species including *P. aeruginosa* (Figure 2A), *Acinetobacter baumannii* (Figure 2B), and *E. coli* (Figure 2C). In addition, we found strong correlation (blue boxes) between the experimental and predicted PAM sequences of 4 species: *S. pyogenes* (Figure 2D), *C. difficile* (Figure 2E), *L. bacterium* (Figure 2F), and *B. halodurans* (Figure 2G). An additional two species had weak correlation (yellow boxes): *S. thermophilus* (Figure 2H) and *T. denticola* (Figure 2I). Weak correlation species are defined here as partial matches where half or less of the predicted PAM matched with the experimentally derived sequence. Finally, we did not find a match (red boxes) for the 3 remaining species: *N. meningitidis* (Figure 2J), *C. jejuni* (Figure 2K), and *S. aureus* (Figure 2L).

**Figure 2.**
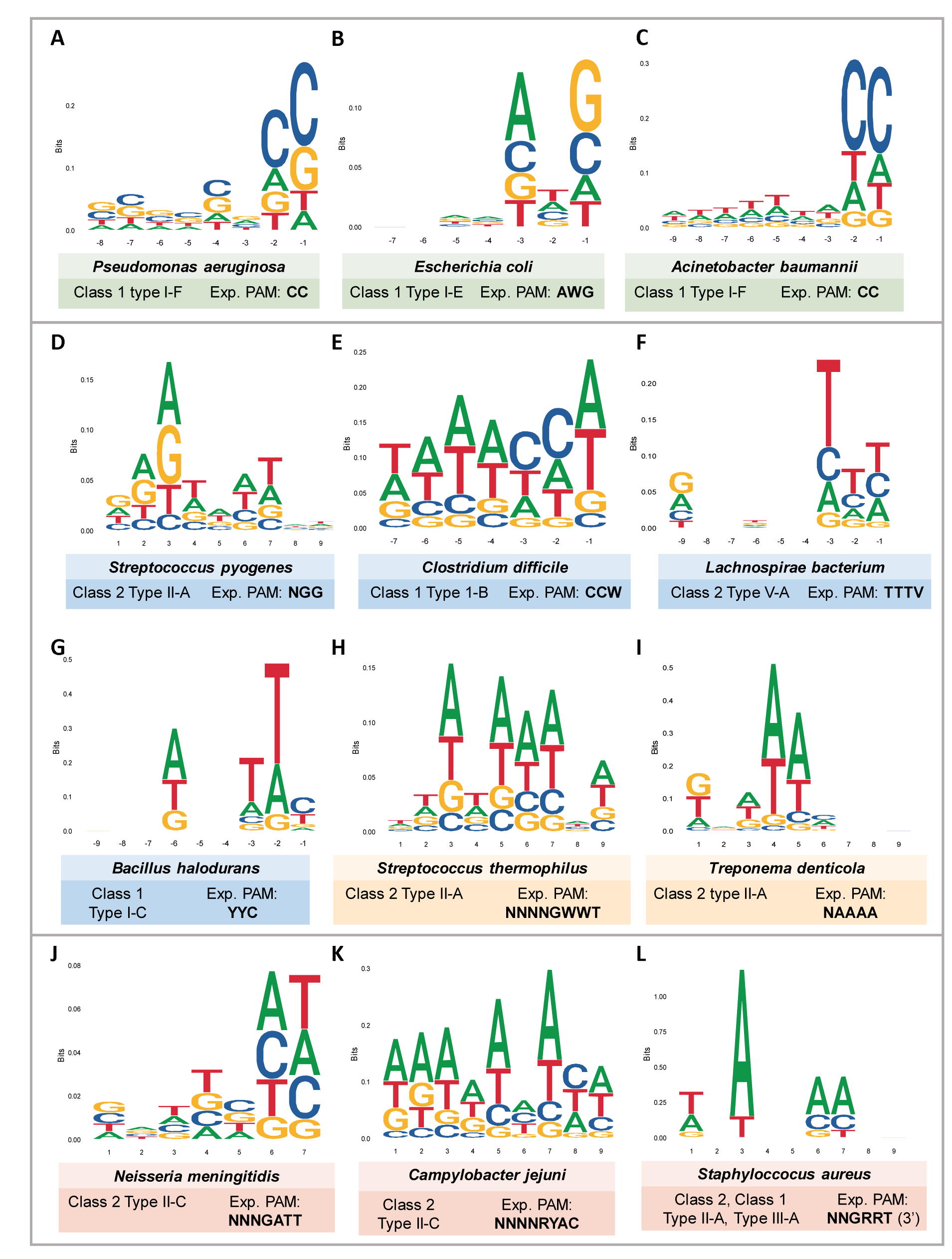
Predicted PAM sequences for experimentally validated species

In our analysis, we observed that the complete and strong partial matches appeared in species that have relatively high spacer hits and consensus Cas-types (Supplementary Table S1). In contrast, the non-matches correlated to a smaller number of hits. A notable exception is *Bacillus halodurans*, which despite a relatively low number of hits returned a strong match to its experimental PAM. (Table 1). Interestingly, type II species seem particularly prone to inaccurate predictions of the PAM sequence, despite large amounts of hits and a consensus Cas-type in certain scenarios. In contrast to type I systems where the PAM sequence may be attributed to either the acquisition or interference stage, type II interference mechanisms have been shown to depend heavily on an accurate PAM sequence [6, 12-14]. Because sequencing results belong to existing phages and other mobile genetic elements, i.e. those that have survived and escaped CRISPR-Cas interference, they may have acquired favorable mutations in the PAM identification region, contributing to an incorrect PAM sequence being derived by CASPERpam.

**Table 1:**
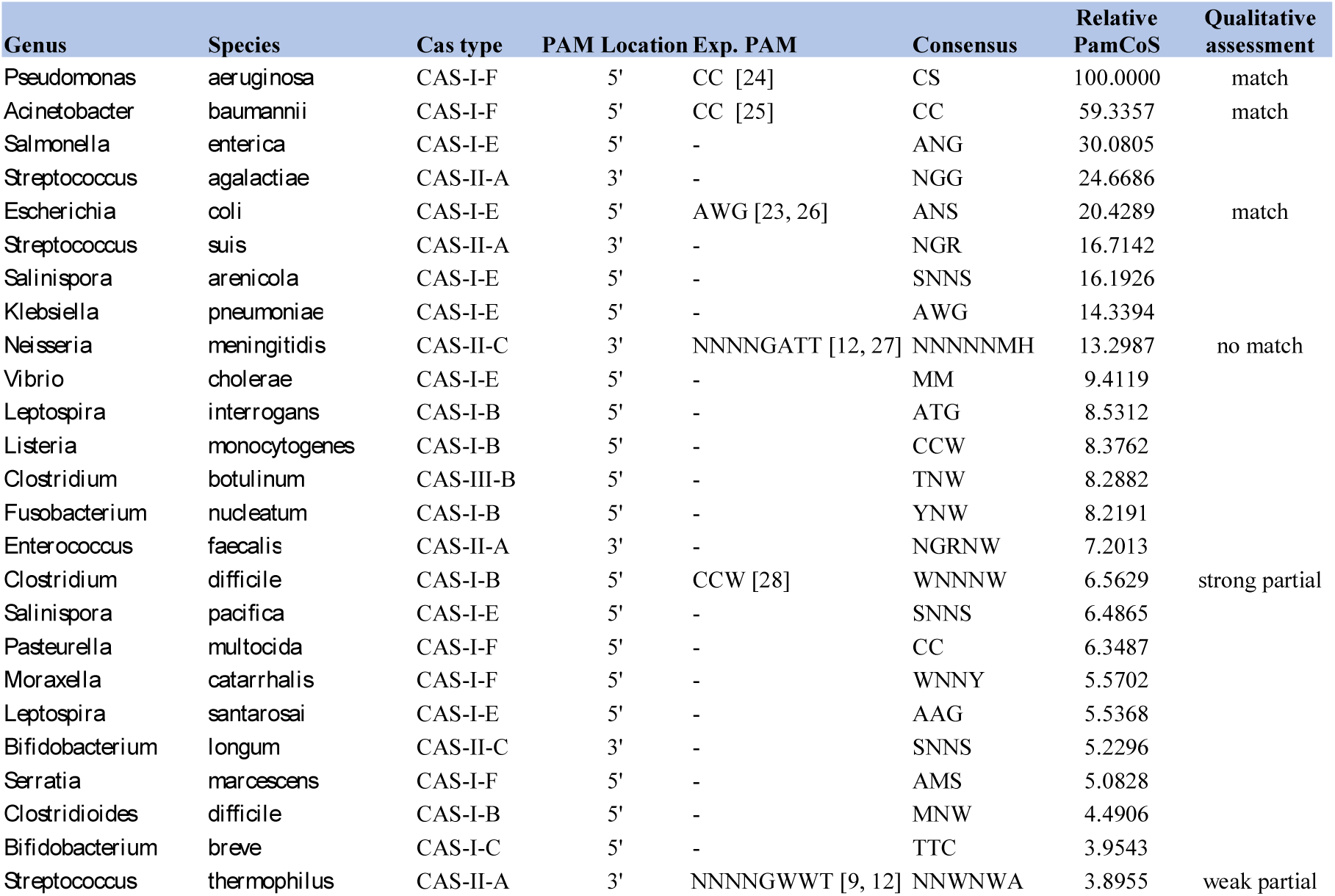
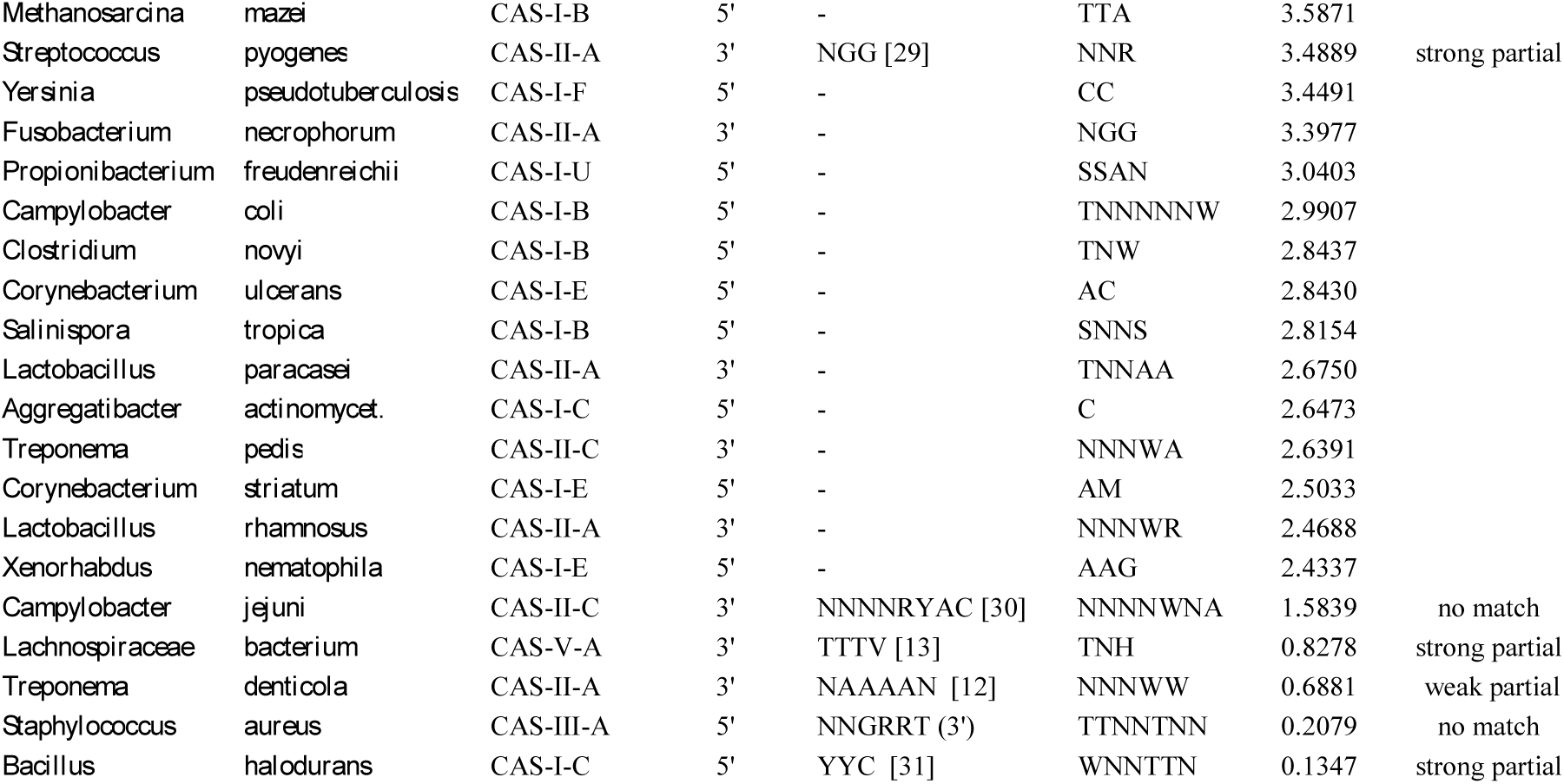
List of select species with top PamCoS scores

Overall, the CASPERpam algorithm is capable of accurately predicting PAMs, particularly for Class 1 systems. The result also highlights the importance of a more quantitative metric to determine whether the information available for a given species is sufficient to predict its putative PAMs, as presented in the next section.

### 3.2. Quantification of the quality of *in silico* PAM prediction

To quantify the accuracy of PAM prediction for a given species, we developed a metric, the PAM confidence score (PamCoS), to determine whether the information obtained from the database (NCBI RefSeq) is of sufficient quantity and quality. PamCoS is calculated as follows:

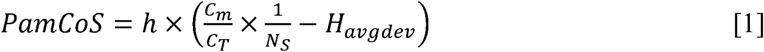

In equation 1, the value *h* represents the number of hits from all the spacers (Supplementary Table S1). C_m_ and C_T_ refer to the number of detected Cas arrays that match the consensus type for a designated species and the total number of arrays, respectively [1]. N_s_ =1 for a species with a well-defined consensus Cas type. However, if a species is not found in the classification database (Supplementary Table S2), the value of N_S_ is increased by 1 for every level up in the phylogenetic tree that the species must be traced until matching a relative. The value H_avgdev_ is the average deviation of the Shannon entropy term for each of the positions deemed as significant contributors to the PAM sequence (denoted by the subscript “sig”).H_avgdev_ is calculated as follows:

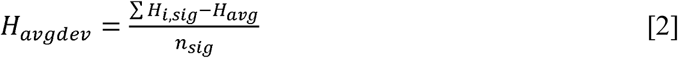

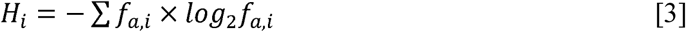

In equation 2, H_i,sig_ is the Shannon entropy of a significant position; H_avg_ represents the average Shannon entropy across all positions in the flanking sequence; and n_sig_ is the number of significant positions. The *f*_*a,i*_ term in equation 3 is the relative frequency of a given nucleotide “a” at position i.

We used PamCoS to score all 1049 species in the CASPERpam produced database (Figure 1 and Supplementary Table S1). The species with the top 40 PamCoS scores are presented in Table 1, along with a select number of additional species for which an experimentally derived PAM sequence is available. The relative PamCoS score is shown in this table, where the scores have been normalized (0 to 100) based on the highest PamCoS score of *Pseudomonas aeruginosa* (2851.61) in the database. High PamCoS scores correlate well with accurate predictions by CASPERpam of the species with experimentally derived PAMs, with the exception of type II species. It does seem however that after a certain PamCoS threshold, the prediction for type II species becomes effective. For instance, *S. agalactiae* and *S. suis*, two species with type II-A CRISPR-Cas systems, have PamCoS scores over 400 (Supplementary Table S1) and exhibit the NGG motif as their predicted PAM, the same as their experimentally proven close relative *S. pyogenes*. This suggests that a PamCoS score of 400 can be used as a minimum threshold to guarantee accurate PAM prediction for all Cas types. While only 8 species in the NCBI RefSeq database currently satisfy this criteria, further sequencing efforts of microbial genomes will aid in providing the necessary information for accurate *in silico* PAM prediction.

To assign a more flexible cutoff for the currently available data and using the PamCoS scores of species with known PAMs, a score above 10 (0.35 relative PamCoS to *P. aeruginosa*) seemed a reasonable threshold for PAM prediction. This however is not a strict cutoff as the prediction of the PAM for *B. halodurans* proves lower PamCoS scoring species may still lead to reasonably accurate PAMs. Lower PamCoS rated species must be viewed with some skepticism, however. Due to the smaller number of hits, even when accompanied by relatively low Shannon entropy values, these predicted sequences are more sensitive to false assignment of significant positions and nucleotides. Further, false sequence noise may be compounded by potential Cas type ambiguity, as a non-consensus Cas-type hit may be incorporated into the PAM sequence prediction.

Overall, use of the PamCoS score enhances *in silico* PAM analysis. Notably, PamCoS allows the PAM sequences produced by CASPERpam to be assessed for their relative accuracy and aid in narrowing the search space of sequences for experimental validation. For instance, species with low PamCoS scores should be validated with a comprehensive *in vitro* or *in vivo* methodology, using the results of CASPERpam as supplemental validation of experimental discovery. On the other hand, species with relatively high PamCoS scores may allow the CASPERpam prediction to be supplemented with only minimal experimentation to provide confirmation of the predicted sequence.

Based on CASPERpam analysis, the number of species with PamCoS scores over the threshold value of 10 (221) represents a very small portion of the tens of thousands of bacterial and archaeal species that have been sequenced. It does however represent an order of magnitude increase in the number of CRISPR-Cas systems that have had a PAM sequence predicted or validated. The result provides a starting point to harness the extensive diversity present across the CRISPR-Cas space.

Of the 221 species above the threshold value, 78% had a Class 1 system, an understudied relative to their more popular single effector Class 2 cousins. Even though the Type I system appears to have a greater variety in their PAM sequences [23], only a select number of nucleotide patterns of PAMs are commonly used. These select PAM motifs are not confined to a single Cas type, rather they are spread across systems (Figure 3A). One potential explanation is that the PAM motif of Class 1 multi-effector systems is likely subject to some degree of horizontal gene transfer. Particularly, the Cas8/10 and Cas1/2 proteins, likely responsible for PAM recognition during the interference and acquisition steps respectively, may be traded within microbial communities, creating within a species hybrid Class 1 systems with a clearly assigned Cas-type using a PAM largely associated with another.

**Figure 3.**
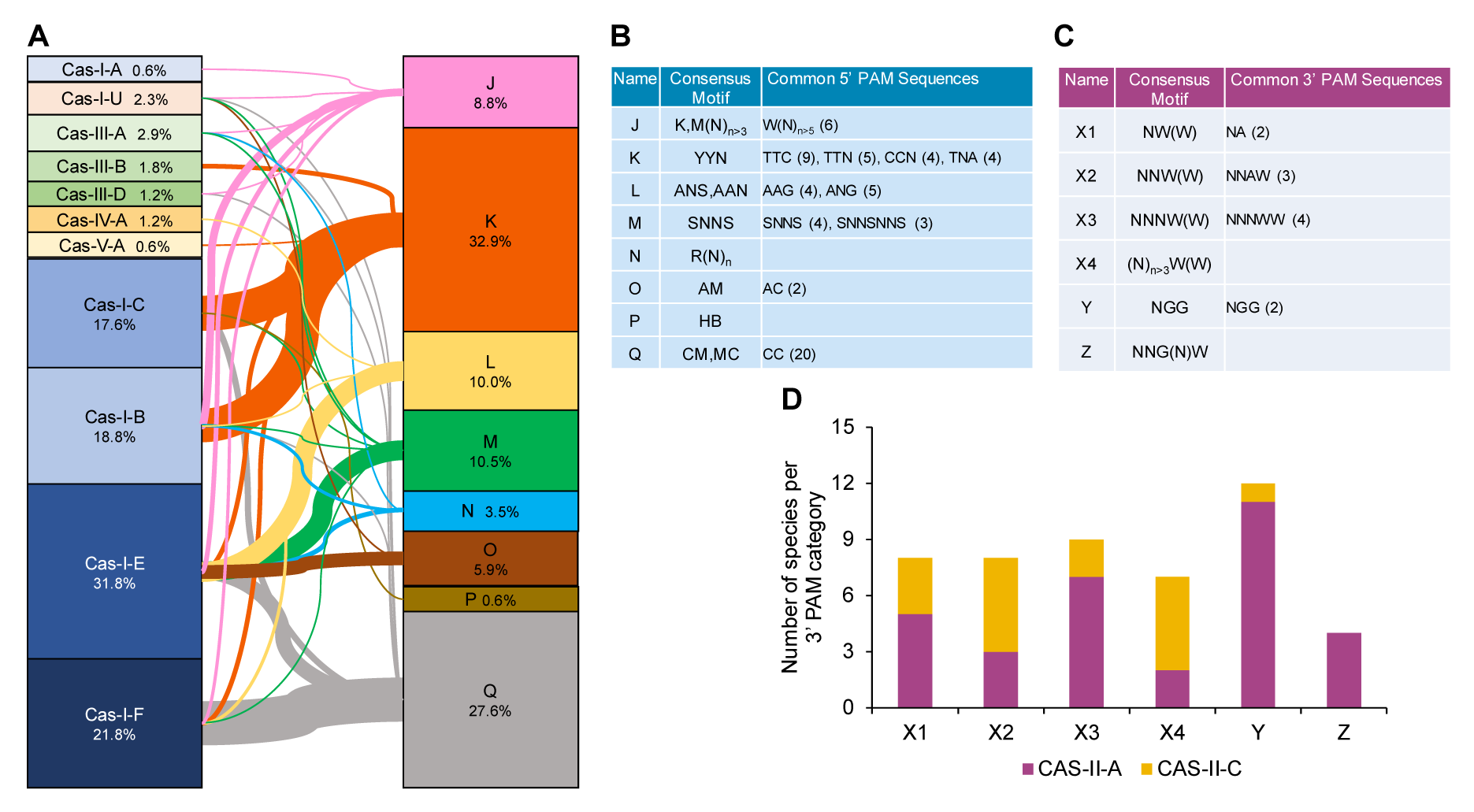
Predicted PAM sequences for select species

### 3.3. PAM prediction highlights diversity and patterns within and between Cas types

To analyze the PAM diversity across different Cas-types, we gathered all PAM motifs of the 221 species above the threshold value and grouped them into families of closely associated motifs. We identified common patterns and assigned 5’ and 3’ PAM sequence categories and/or subcategories to these sets for 218 species (Figures 3B and 3C). Each category/subcategory is defined by a consensus motif that captures the general pattern. Three species from the original set of 221 species, including *Proteus mirabilis, Bifidobacterium longum,* and the *Eubacterium* sp., were removed from the analysis because CASPERpam could not assign either a consensus Cas-type or the PAM motif was clearly the result of multiple Cas-types.

The 5’ PAM sequences contain 8 categories (J, K, L, M, N, O, P, and Q) (Figure 3A, 3B). The K and Q categories represent 61% of PAMs located on the 5’ end. The ‘CC’ PAM of the Q category and ‘TTC’ PAM of K are heavily represented (Figure 3A, 3B). These groups are disproportionately associated with types I-F and I-B respectively; however, they are not exclusive and also appear in species with other Cas-types. The PAM sequence space of the 170 species with reliable 5’ PAMs (as defined by the confidence score) can be seen mapped together in Figure 3A.

It is striking to observe that none of the Cas types nor subtypes map to a unique PAM category, with the exception of a few types with less than 5 species represented in the sample (e.g. Cas-III-B and Cas-I-A). This result supports the view that the current CRISPR-Cas classification method is imperfect likely due to the CRISPR-Cas systems’ similarity and modularity prone to horizontal gene transfer [1]. It also prevents a simple classification based on sequence homology or phylogenetics from perfectly partitioning the heterogeneity of these systems.

The 3’ PAM sequences consists of 3 categories (X, Y, and Z) with the X category broken into a further 4 subcategories (X1, X2, X3, X4) (Figure 3C). Among the 3’ PAM sequences, the category Y, with the NGG consensus motif, is quite common, representing 25% of identified PAMs (Figures 3C, 3D). The category X, although representing two-thirds of identified PAMs, has a much more broadly defined consensus motif (N)_n_W and lent itself to subcategorization of which none reached a representation comparable to the Y category (Figures 3C, 3D). The ‘NGG’ PAM found in the Y category is the most represented PAM motif, but CASPERpam only identified two such species, with an exact ‘NGG’ PAM. Other PAMs in the category were defined more broadly, with adenines and sometimes thymines also highly represented at the 2 and 3 positions creating an NRR, and NKK PAMs. This result is consistent with experimental evidence that type II-A systems such as the canonical *S. pyogenes* exhibited some affinity for a ‘NAG’ PAM. Even though this detected promiscuity may be overstated by the *in silico* analysis, it highlights the importance of future experimental studies on species with a PAM falling in the Y category to focus on the relative affinity for guanine over adenine or thymine.

Based on the 5’ PAM space and the X and Z 3’ PAM categories, we clearly observed that there exists a high level of non-specificity in many PAM interacting domains. It cannot be discounted that this diversity may be underrepresented by the *in silico* technique, as only those with particularly effective targeting will appear across the targeted genomes [5]. On the other hand, PAM sequences are likely to be mutated for phages to survive, and thus sequencing assemblies may not represent the true form of the PAM sequence, but rather an overrepresentation of a random set of mutated sequences despite a close spacer match. Such a hypothesis would result in the *in silico* process revealing greater non-specificity than would be shown experimentally. Likely, these two effects are both present, and result in some overly specific as well as some overly promiscuous sequence predictions from the *in silico* method.

## 4. Conclusion

To harness a large, diverse class of alternative CRISPR systems outside of the group of well studied Cas9 and Cas12 variants, it is critical to identify PAM sequences for characterizing these novel CRISPR systems. Development of CASPERpam provides a powerful tool to predict PAMs for a given species and allows subsequent experiments to change from a determination to a validation study. While some previous studies have performed such an *in silico* PAM analysis for a select number of species to guide experimental studies of native CRISPR systems, this study has performed an exhaustive PAM analysis for every CRISPR-Cas carrying bacterial and archaeal genome in the NCBI database. As research moves into investigating a greater number of CRISPR-Cas systems with novel functionalities, CASPERpam will be a useful bioinformatic tool for those seeking to understand and harness the diversity of CRISPR systems.

## Abbreviations

CRISPR: Clustered Regularly Interspaced Palindrome Repeats
CASPER: CRISPR Associated Software for Pathway Engineering and Research
PAM: Protospacer Adjacent Motif
PamCoS: PAM confidence score
NCBI: National Center for Biotechnology Information

## Acknowledgements

The research is funded by the DARPA YFA award #D17AP00023. The views, opinions, and/or findings contained in this article are those of the authors and should not be interpreted as representing the official views or policies, either expressed or implied, of the funding agencies.

## References

1. Makarova, K.S., et al., An updated evolutionary classification of CRISPR-Cas systems. Nat Rev Micro, 2015. 13(11): p. 722–736.

2. Barrangou, R., Diversity of CRISPR-Cas immune systems and molecular machines. Genome Biol, 2015. 16: p. 247.

3. Shah, S.A., et al., Protospacer recognition motifs: Mixed identities and functional diversity. RNA Biology, 2013. 10(5): p. 891–899.

4. Mojica, F.J.M., et al., Short motif sequences determine the targets of the prokaryotic CRISPR defence system. Microbiology, 2009. 155(3): p. 733–740.

5. Leenay, R.T. and C.L. Beisel, Deciphering, Communicating, and Engineering the CRISPR PAM. Journal of Molecular Biology, 2017. 429(2): p. 177–191.

6. Hsu, P.D., et al., DNA targeting specificity of RNA-guided Cas9 nucleases. Nat Biotechnol, 2013. 31(9): p. 827–32.

7. Makarova, K.S. and E.V. Koonin, Annotation and Classification of CRISPR-Cas Systems, In CRISPR: Methods and Protocols, M. Lundgren, E. Charpentier, and P.C. Fineran, Editors. 2015, Springer New York: New York, NY. p. 47–75.

8. Cebrian-Serrano, A. and B. Davies, CRISPR-Cas orthologues and variants: optimizing the repertoire, specificity and delivery of genome engineering tools. Mamm Genome, 2017. 28(7-8): p. 247–261.

9. Horvath, P., et al., Diversity, activity, and evolution of CRISPR loci in Streptococcus thermophilus. J Bacteriol, 2008. 190(4): p. 1401–12.

10. Ran, F.A., et al., In vivo genome editing using Staphylococcus aureus Cas9. Nature, 2015. 520(7546): p. 186–91.

11. Elmore, J., et al., DNA targeting by the type I-G and type I-A CRISPR–Cas systems of Pyrococcus furiosus. Nucleic Acids Research, 2015. 43(21): p. 10353–10363.

12. Esvelt, K.M., et al., Orthogonal Cas9 proteins for RNA-guided gene regulation and editing. Nature methods, 2013. 10: p. 1116–21.

13. Zetsche, B., et al., Cpf1 is a single RNA-guided endonuclease of a class 2 CRISPR-Cas system. Cell, 2015. 163(3): p. 759–71.

14. Karvelis, T., et al., Rapid characterization of CRISPR-Cas9 protospacer adjacent motif sequence elements. Genome Biology, 2015. 16: p. 253.

15. Biswas, A., et al., CRISPRTarget: Bioinformatic prediction and analysis of crRNA targets. RNA Biology, 2013. 10(5): p. 817–827.

16. Pyne, M.E., et al., Harnessing heterologous and endogenous CRISPR-Cas machineries for efficient markerless genome editing in Clostridium. 2016. 6: p. 25666.

17. Mendoza, B.J. and C.T. Trinh, Enhanced guide-RNA design and targeting analysis for precise CRISPR genome editing of single and consortia of industrially relevant and non-model organisms. Bioinformatics, 2018. 34(1): p. 16–23.

18. Shmakov, S.A., et al., The CRISPR Spacer Space Is Dominated by Sequences from Species-Specific Mobilomes. mBio, 2017. 8(5): p. e01397–17.

19. Xue, C., et al., CRISPR interference and priming varies with individual spacer sequences. Nucleic Acids Research, 2015. 43(22): p. 10831–10847.

20. Westra, Edze R., et al., CRISPR Immunity Relies on the Consecutive Binding and Degradation of Negatively Supercoiled Invader DNA by Cascade and Cas3. Molecular Cell. 46(5): p. 595–605.

21. van der Oost, J., et al., Unravelling the structural and mechanistic basis of CRISPR–Cas systems. Nature Reviews Microbiology, 2014. 12: p. 479.

22. Wagih, O., ggseqlogo: a versatile R package for drawing sequence logos. Bioinformatics, 2017. 33(22): p. 3645–3647.

23. Fu, B.X., et al., High-Throughput Characterization of Cascade type I-E CRISPR Guide Efficacy Reveals Unexpected PAM Diversity and Target Sequence P. Genetics, 2017. 206(4): p. 1727–1738.

24. Cady, K.C., et al., The CRISPR/Cas adaptive immune system of Pseudomonas aeruginosa mediates resistance to naturally occurring and engineered phages. J Bacteriol, 2012. 194(21): p. 5728–38.

25. Karah, N., et al., CRISPR-cas Subtype I-Fb in Acinetobacter baumannii: Evolution and Utilization for Strain Subtyping. PLOS ONE, 2015. 10(2): p. e0118205.

26. Luo, M.L., et al., Repurposing endogenous type I CRISPR-Cas systems for programmable gene repression. Nucleic Acids Research, 2015. 43(1): p. 674–681.

27. Hou, Z., et al., Efficient genome engineering in human pluripotent stem cells using Cas9 from <Neisseria meningitidis>. Proceedings of the National Academy of Sciences, 2013. 110(39): p. 15644.

28. Boudry, P., et al., Function of the CRISPR-Cas System of the Human Pathogen Clostridium difficile. MBio, 2015. 6(5): p. e01112–15.

29. DiCarlo, J.E., et al., Genome engineering in Saccharomyces cerevisiae using CRISPR-Cas systems. Nucleic Acids Res, 2013. 41(7): p. 4336–43.

30. Kim, E., et al., In vivo genome editing with a small Cas9 orthologue derived from Campylobacter jejuni. Nature Communications, 2017. 8: p. 14500.

31. Leenay, Ryan T., et al., Identifying and Visualizing Functional PAM Diversity across CRISPR-Cas Systems. Molecular Cell, 2016. 62(1): p. 137–147.

